# Loss of *Slc35a2* alters development of the mouse cerebral cortex

**DOI:** 10.1101/2023.11.29.569243

**Authors:** Soad Elziny, Sahibjot Sran, Hyojung Yoon, Rachel R. Corrigan, John Page, Amanda Ringland, Anna Lanier, Sara Lapidus, James Foreman, Erin L. Heinzen, Philip Iffland, Peter B. Crino, Tracy A. Bedrosian

**Author notes:** equal contributors (Soad Elziny, Sahibjot Sran, Peter B. Crino, Tracy A. Bedrosian). Correspondence to: Tracy A. Bedrosian, Ph.D. Nationwide Children’s Hospital Research Building IV 575 Children’s Crossroad Columbus, OH 43215.

## Abstract

Brain somatic variants in *SLC35A2* are associated with clinically drug-resistant epilepsy and developmental brain malformations, including mild malformation of cortical development with oligodendroglial hyperplasia in epilepsy (MOGHE). *SLC35A2* encodes a uridine diphosphate galactose translocator that is essential for protein glycosylation; however, the neurodevelopmental mechanisms by which *SLC35A2* disruption leads to clinical and histopathological features remain unspecified. We hypothesized that focal knockout (KO) or knockdown (KD) of *Slc35a2* in the developing mouse cortex would disrupt cerebral cortical development through altered neuronal migration and cause changes in network excitability. We used *in utero* electroporation (IUE) to introduce CRISPR/Cas9 and targeted guide RNAs or short-hairpin RNAs to achieve *Slc35a2* KO or KD, respectively, during early corticogenesis. Following *Slc35a2* KO or KD, we observed disrupted radial migration of transfected neurons evidenced by heterotopic cells located in lower cortical layers and in the sub-cortical white matter. *Slc35a2* KO in neurons did not induce changes in oligodendrocyte number, suggesting that the oligodendroglial hyperplasia observed in MOGHE originates from distinct cell autonomous effects. Spontaneous seizures were not observed, but intracranial EEG recordings after focal KO showed a reduced seizure threshold following pentylenetetrazol injection. These results demonstrate that *Slc35a2* KO or KD *in vivo* disrupts corticogenesis through altered neuronal migration.

## Introduction

*SLC35A2* is an X-linked gene that encodes a UDP-galactose transporter responsible for sequestering galactose within the Golgi lumen for post-translational modification of proteins. Impairing SLC35A2 function, and thereby depleting the luminal supply of galactose, disrupts glycosylation machinery and leads to defective modification of proteins and lipids^1^. Protein glycosylation is critical to diverse aspects of neural development and function, including cellular adhesion and migration, synaptic transmission, and membrane excitability^2^. Germline variants in *SLC35A2* cause congenital disorders of glycosylation (*SLC35A2*-CDG)^3^, which are highly linked to severe seizures (i.e., epileptic encephalopathy) and other organ system phenotypes^4^.

Interestingly, brain somatic *SLC35A2* variants are associated with malformations of cortical development (MCD), which represent the most common histopathologic diagnosis in pediatric patients with drug resistant epilepsy^5^. *SLC35A2* somatic variants were initially reported in cases of focal cortical dysplasia (FCD) type I and mild malformation of cortical development (mMCD)^6^ at variant allele fractions as low as 1.9% in surgically resected brain specimens^7,8^. More recently, a novel histopathological entity known as mild malformation of cortical development with oligodendroglial hyperplasia in epilepsy (MOGHE) has been associated with somatic *SLC35A2* variants^9,10^. MOGHE is characterized by oligodendroglial hyperplasia and patchy hypomyelination, as well as persistent embryonic cortical microcolumns and neuronal heterotopia, suggesting a role for *SLC35A2* in assembly of the cortex and integrity of oligodendrocyte maturation during corticogenesis^6,11,12^. Indeed, the most recent revision of classification guidelines on FCD from the International League Against Epilepsy (ILAE) recommended MOGHE as the definitive histopathologic MCD diagnosis associated with *SLC35A2* mutations^13^.

While somatic *SLC35A2* variants are associated clinically with MCD, the causal effects of SLC35A2 loss on cortical development have not been experimentally assessed. We hypothesized that targeted knockout (KO) of *Slc35a2* in the fetal mouse brain would impair cortical development and alter network integrity. Because some *Slc35a2* variants observed in patients may retain partial functionality of the protein, we hypothesized that *Slc35a2* knockdown (KD) would also be sufficient to disrupt cortical development by impairing neuronal migration. To test our hypothesis, we used *in utero* electroporation (IUE) to induce either *Slc35a2* KO or KD in a limited set of neural precursor cells, thus providing a model of human somatic mutations.

## Materials and Methods

### Animals

CD-1 mice (Charles River Laboratories) were housed under a standard light/dark cycle with access to food and water *ad libitum*. All experiments were performed in compliance with the animal care and use guidelines issued by the National Institutes of Health and under protocols approved by the Institutional Animal Care and Use Committees at Nationwide Children’s Hospital and University of Maryland, Baltimore.

### CRISPR/Cas9 construct generation and validation

Guide RNAs (gRNAs) targeting mouse *Slc35a2* exons 2 and 3 were designed and created *in silico* using CHOPCHOP software (chopchop.cbu.uib.no). (gRNA Target sequence A: TGCGAGCGTAGCGGATGCTGAGG, gRNA target sequence B: GGCTGTCCTGGTCCAATATGTGG).

Using the Golden Gate Assembly protocol, *Slc35a2* gRNAs (under a U6 promotor) were spliced into an pX458v2-R-spCas9 plasmid containing an mCherry (*in vitro* experiments) or EGFP fluorescent reporter (*in vivo* experiments). gRNA and plasmid sequences were confirmed by Sanger sequencing (Azenta Life Sciences Co). A scrambled gRNA sequence (confirmed by sequencing) was used as a transfection control.

### shRNA constructs

Plasmids were designed and produced using services offered by VectorBuilder. Each plasmid contained a coding region for a short hairpin RNA (shRNA) and enhanced green fluorescent protein (EGFP) under the control of a U6 and CAG promoter, respectively. Three shRNA sequences were used: shRNA #1 (TTCCACCTGGACCCATTATTTCTCGAGAAATAATGGGTCCAGGTGGAA), shRNA #2 (CTTGCAGAATAACCTCCAGTACTCGAGTACTGGAGGTTATTCTGCAAG), and a scrambled shRNA sequence (CCTAAGGTTAAGTCGCCCTCGCTCGAGCGAGGGCGACTTAACCTTAGG) that targeted no gene in the mouse genome as a control.

### *Slc35a2* knockout cell lines

Neuro2a cells (Sigma-Aldrich, SCC297) were transfected with dual gRNA CRISPR/Cas9-plasmid using Lipofectamine LTX with Plus reagent (Thermo Fisher Scientific, 15338100) and 0% serum EMEM media (QBI-112-018-101) for 48 h. Transfected Neuro2a cells were assayed by flow cytometry (University of Maryland School of Medicine Flow Cytometry Core) and sorted based on mCherry fluorescence. mCherry+ sorted cells were re-plated in complete media and grown to confluence to assess success of *Slc35a2* KO.

*Slc35a2* knockdown cell lines

NIH/3T3 cells (ATCC, CRL-1658) were cultured at 37C (5% CO_2_) in DMEM (Fisher Scientific, 11320082) with 10% Fetal Bovine Serum (Fisher Scientific, 11-648-647) and Antibiotic-antimycotic (Abcam, 15-240-062). Cells were seeded onto T-75cm^2^ flasks (Fisher, FB012937) to reach 75% confluence. DNA plasmid electroporation was performed with NEON Transfection System (Invitrogen, MPK5000) following the manufacturer’s recommendations. Briefly, cultured cells were collected at 70% confluence using two washes with DPBS (Gibco, 14190250) followed by treatment with TrypLE Express Enzyme (Gibco, 12604021) for 3 minutes. Detached cells were collected into 15mL tubes and collected with centrifugation at 400xg for 4 minutes. The resulting pellet was resuspended in DMEM/F-12 and cell density was determined using the Countess 3 Automated Cell Counter (Invitrogen, AMQAF2000) using Trypan blue solution (Gibco, 15250061). Cells were re-pelleted for resuspension in Buffer R (Invitrogen, MPK10025) containing 500 ng of plasmid to a final density of 1x106 cells/mL. Electroporation was carried out at 1400V, for 3 cycles with 2 ms intervals. Cells were then seeded on 24-well plates, cultured with full growth media described above for 2 days, and pelleted for qPCR preparation.

### *Slc35a2* knockout RT-qPCR

Changes in *Slc35a2* mRNA expression after CRISPR KO were assayed by RT-qPCR. Total RNA was extracted from Neuro2A cell pellets of transfected *Slc35a2* KO lines, scramble, and wild-type cells using an RNeasy Mini Kit (Qiagen, 74104). RNA was converted to cDNA using a high-capacity cDNA reverse transcription kit. The following RT-qPCR primers within *Slc35a2* edited regions were used: F: GCCTCTGACTCCTTCCTCCTAT; R: AGGTGAGACCTTTGAGCACTTC). Mouse GAPDH was used as an expression control (GAPDH: F: AGGTCGGTGTGAACGGATTTG; R:TGTAGACCATGTAGTTGAGGTCA). RT-qPCR was performed using SYBR Power Green PCR master mix with a Viia7 qPCR machine (Thermo Fisher). PCR conditions were set to the following: 95°C for 10 min, followed by 45 cycles of 15 s denaturation at 95°C, 1 min annealing at 60°C, followed by 30 s extension at 72°C. Expression of *Slc35a2* was determined relative to a standard curve created using wild-type mouse cDNA at the following concentrations: 1:20, 1:60, 1:180, 1:540 and 1:1620. Negative controls used were DNAse-free water and diluted random primers.

### *Slc35a2* knockdown RT-qPCR

RNA was extracted from cell pellets with Trizol (Invitrogen, 15596018) following manufacturer recommendations. RNA yield was quantified using a NanoDrop (Thermo Scientific ND-2000C) or Qubit (Invtrogen, Q33238). cDNA was generated from 40 ng of extracted RNA using a high-capacity reverse transcription kit (Applied Biosystems, 4368814) and RT-qPCR was performed on the QuantStudio Pro 6 system (Thermo Scientific, A43180) using PowerTrack SYBR Green Master Mix (Applied Biosystems, A46109). Pre-designed primers for *Slc35a2* (Mm.PT.58.29074836) and *Gapdh* (Mm.PT.39a.1) were obtained from Integrated DNA Technologies. Temperature and cycling parameters were: 2 minutes at 50°C, 10 minutes at 95°C, and 40 cycles of 15 seconds at 95°C and 60 seconds at 60°C. Relative expression of *Slc35a2* was determined using the delta delta Ct method.

### *Slc35a2* knockout western assay

For CRISPR experiments, Neuro2a cells were lysed in radioimmunoprecipitation assay (RIPA) buffer with protease and phosphatase inhibitors. 30 µg of protein lysate supernatants were run on a Bolt BT Plus 4–12% gel (Invitrogen) and transferred onto PVDF membranes at 4°C. Membranes were probed overnight at 4°C with antibodies for SLC35A2 (Sigma Aldrich HPA036087 Rb 1:100, polyclonal) and GAPDH (Abcam Mouse 1:1000; monoclonal). Densitometry analysis (Origin software) was performed on blots with technical replicates in triplicate.

### *Slc35a2* developmental expression

Cerebral hemispheres from wild-type CD-1 mice were collected and snap frozen for protein analysis at embryonic day 16 and post-natal days 0, 2, 7, and 14. To prepare protein homogenates, tissue was rapidly weighed and homogenized with a pestle homogenizer in the following solution: 1x Cell Lysis Buffer (Cell Signaling, 9803S), 0.5% (v/v) Dithiothreitol (ThermoFisher Scientific, 327190010), 1% (v/v) PMSF (Cell Signaling, 8553), 2% (v/v) Phosphatase Inhibitor Cocktail (Millipore Sigma, 524629), and 0.5% (v/v) Phosphatase Inhibitor Cocktail Set V (Millipore Sigma, 524629). The resulting lysates were centrifuged to collect supernatant for downstream Western Blotting. Protein concentrations were determined via Rapid Gold BCA Assay (Thermo Scientific, A53225). Protein samples were incubated at 37°C for 1 hour. 15 µg of total protein was resolved by sodium dodecyl sulfate-polyacrylamide gel electrophoresis (SDS-PAGE) using a 10-20% tris-glycine gradient (Invitrogen, XP10205BOX), transferred to a PVDF membrane using the Trans-Blot Turbo Transfer System (BioRad, 1704150), and blocked in 10% milk for an hour. Membranes were probed with an SLC35A2 polyclonal antibody overnight at 4°C (1:2000, Invitrogen, PA5-77178), followed by incubation with goat anti-rabbit HRP-linked secondary antibody (1:1000, Cell Signaling, #7074) for 1 hour at room temperature. Beta-tubulin (1:10000, Abcam, ab6046) with secondary antibody (1:1000, Cell Signaling, #7076) was used as a loading control. Membranes were developed with a chemiluminescent HRP substrate (Millipore Sigma, WBKLS) and imaged using a ChemiDoc MP (BioRad). SLC35A2 protein expression was quantified and normalized to the loading control using ImageJ software (NIH).

### *In utero* electroporation

CD-1 dams were anesthetized with isoflurane at embryonic day 14 (E14) for KO experiments or embryonic day 15.5 (E15.5) for KD experiments and the uterine horns were surgically exposed as described previously^14,15^. Neuroglial progenitor cells transfected at E14-E15.5 are destined to become layer II/III cortical neurons^16^. Three to seven embryos per dam were injected with plasmid DNA (approximately 1 µg/µl) in the lateral ventricle using a pulled glass micropipette. Embryos were electroporated with a BTX ECM830 square wave pulse generator (5 electrical pulses at 50V with a 50msec pulse duration and 500 msec pulse interval) by holding the embryo head in between the electrode forceps with the anode oriented apically to direct the plasmid toward neural progenitor cells lining the lateral ventricle. Buprenorphine was administered subcutaneously to the dam as an analgesic following the procedure.

At postnatal day 1 (P1), pups were examined for EGFP fluorescence through the skull using a NIGHTSEA Xite EGFP Fluorescence Flashlight to assess positive transfection. Pups exhibiting EGFP fluorescence (CRISPR KO-, shRNA KD-, or controls) were selected for immunohistochemical analysis and euthanized under deep anesthesia at P1, P4, or P10 timepoints depending on the experiment. All mice to be processed for immunohistochemical analysis underwent transcardial perfusion with 4% paraformaldehyde in 0.1M phosphate buffer and brains were extracted and prepared for histological analysis.

### Immunofluorescence

For KO experiments, whole brains were post-fixed in 4% paraformaldehyde overnight, paraffin embedded into cassettes, and microtome sectioned at 8 µm on a Dolbey-Jamison as325. Sections were mounted on Superfrost Plus Gold slides (Fisher Scientific) and allowed to dry overnight before proceeding with immunostaining. Sections were deparaffinized in xylenes and graded alcohols, then probed with various antibodies or Cresyl violet before cover slipping with toluene. Sections were probed with Ctip2 (1:100; rat monoclonal; Abcam ab18465), EGFP (1:500; chicken polyclonal; Abcam ab13970), and SATB2 (1:500; mouse monoclonal; Abcam ab51502), or EGFP and Olig2 (1:500; rabbit monoclonal; Abcam ab109186) primary antibodies. The following were used for visualization: Alexa 488 A-11039 (EGFP), 647 A-21247 (Ctip2) and 594 A-21203 (SATB2) or 3,3′-Diaminobenzidine (DAB, SIG-D5637-1G) (Olig2). Immunofluorescence sections were counterstained with DAPI (Fluoroshield with DAPI, Sigma Aldrich, F6057) and mounted using Prolong Gold Antifade Mountant (Invitrogen, P36930).

For shRNA experiments, brains were post-fixed in 4% paraformaldehyde solution for 24 hours, then moved to a 30% sucrose solution before sectioning with a sliding microtome (Leica SM2010R) at a thickness of 30 µm. Sections were washed with 0.1M phosphate buffered saline (PBS), counterstained with DAPI (1:1000, BioLegend, 422801) and mounted on glass slides using Prolong Gold Antifade Mountant (Invitrogen, P36930).

### Image analysis

One to three brain sections from each animal were selected for image analysis based on the region of greatest extent of electroporation, i.e., EGFP fluorescence. Sections from KO experiments were analyzed as images taken on a spinning-disk W1 confocal microscope and reconstructed in Imaris Bitplane software. To determine the effects of *Slc35a2* KO, the number of cells expressing EGFP in cortical layers I–VI and the corpus callosum was quantified and expressed relative to the total number of EGFP-positive cells. To determine the proportion of malpositioned neurons within *Slc35a2* KO brains in comparison to EGFP control, the number of positively transfected neurons was manually counted and binned for each brain (n=5 brains per group). Bins were either layer II/III, IV-VI, or the subcortical white matter, with the layer II/III being the appropriate laminar destination and neurons in both the layer IV-VI bin or the subcortical white matter being classified as malpositioned. EGFP-positive neurons were deemed malpositioned if they were observed outside of the expected cortical layers II/III according to birthdate at the time of electroporation (E14). Olig2-positive cells in each section were manually counted for the entire section. For KD experiments, images were acquired using a Zeiss Axio Imager 2 (Carl Zeiss AG) and subsequently imported into ImageJ for analysis. Six regions of interest (ROI) from the ventricular zone to the outer cortex were defined according to DAPI staining. The total area of EGFP present in each ROI was calculated and expressed as a percentage of the sum of the EGFP area in all ROIs.

### *In vivo* EEG recordings

*Slc35a2* KO mice, EGFP-transfected control mice, and wild-type littermates underwent EEG electrode implantation surgery, EEG recording, and PTZ testing at P60 or P120. Briefly, EEG electrodes were implanted and secured onto the skull 3 mm behind bregma (and positioned evenly across the midline) under isoflurane anesthesia. Continuous EEG recordings (72 h) were performed using the Pinnacle Technology 3 channel EEG system (Lawrence). Electroporated *Slc35a2* KO, EGFP control mice, or wild-type mice were administered the pro-convulsant agent pentylenetetrazol (PTZ) at 55 mg/kg intraperitoneally and recorded for 40 min to capture the time to first electrographic seizure detected by EEG. PTZ reliably evokes convulsive seizures that are characterized according to the Racine scale in mice^17^. All mice were video recorded and electrographically recorded for 10 minutes as a baseline, then administered PTZ and scored according to the Racine scale every 2 minutes for 30 minutes. A seizure was defined electrographically as 10 seconds or more of rhythmic, continuous, sharply contoured, and evolved activity with a clear onset and clear offset. The behavioral correlate of this activity was tonic-clonic activity (Racine score of 4-5^18^). Thirty minutes after PTZ injection the animals were euthanized using CO_2_ asphyxiation followed by cervical dislocation. Behavioral and electrographic seizures were confirmed by reviewing recordings blinded to condition after termination of the experiment.

### Statistical analysis

The proportion of malpositioned neurons to appropriately positioned neurons, as well as Olig2-positive cell density, in KO experiments was compared using Students unpaired t-test, accepting p<0.05 as level of significance. Seizure latency was compared using a Student’s unpaired t-test (p<0.05) with Welch’s correction between the following groups at P60 and P120: EGFP vs WT, EGFP vs KO, EGFP vs KO. Comparisons of KD neurons per cortical bin were analyzed with two-way ANOVA followed by Tukey’s post-hoc comparisons. All statistical analysis was done using Graphpad Prism software.

## Results

### SLC35A2 expression in cerebral cortex

In mice, corticogenesis occurs between embryonic days (E) 10.5 to 18.5. We quantified SLC35A2 protein expression in the mouse forebrain at E14, 16, and postnatal days (P) 0, 2, 7, and 14 by Western blot to determine baseline expression. SLC35A2 was highly expressed at embryonic and postnatal time points (Fig. 1a).

**Fig. 1.**
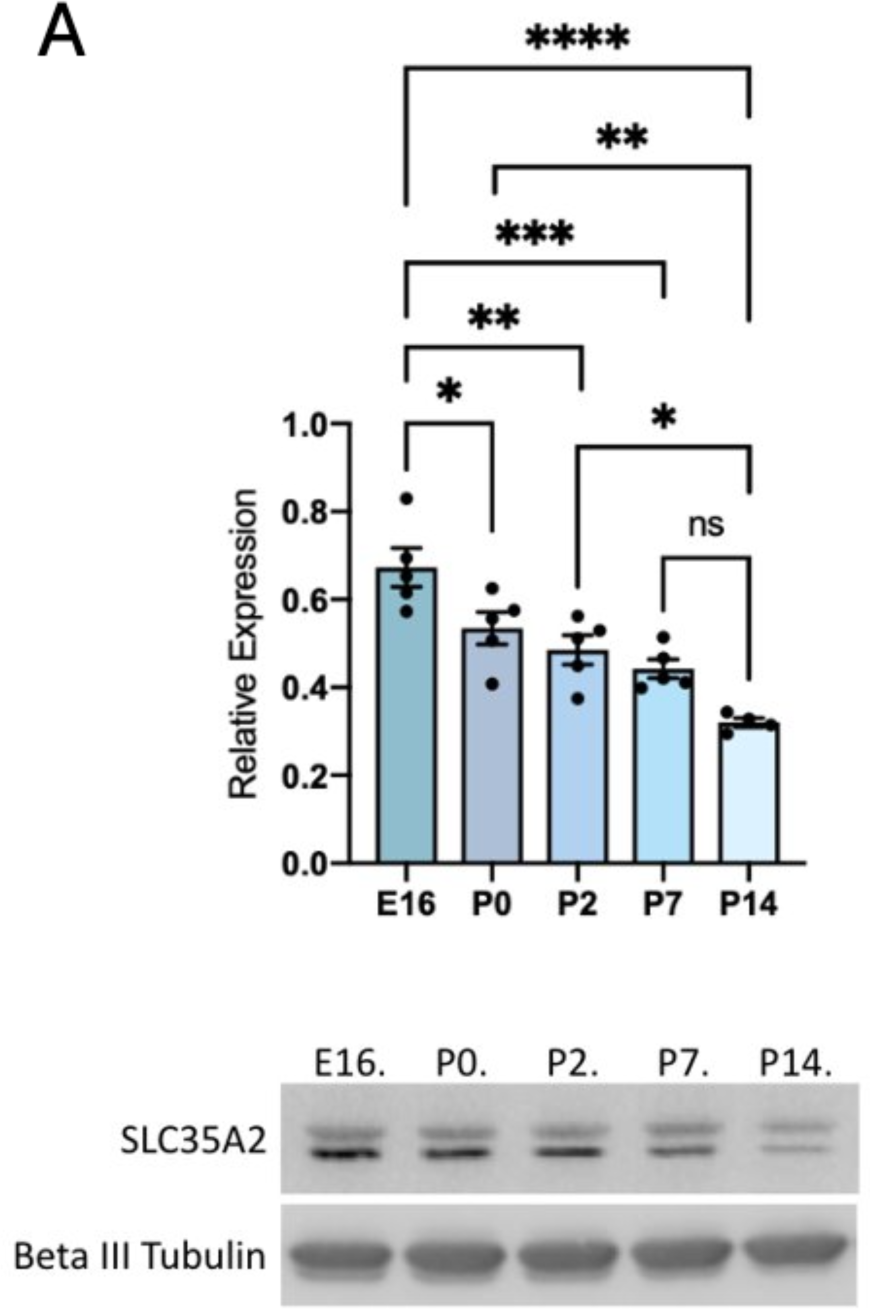
SLC35A2 is highly expressed in the mouse cortex during development. A) Western Blot of SLC35A2 expression relative to GAPDH depicting expression in mouse cortex across development. Both SLC35A2 isoforms were quantified together. E = embryonic day, P = postnatal day. Mean ± SEM. * P<0.05, ** P<0.01, *** P<0.001.

### Validation of *Slc35a2* KO *in vitro*

To knockout *Slc35a2* expression, we designed a CRISPR/Cas9 strategy targeting exons 2 and 3 (Fig. 2a). To confirm the knockout efficiency, we transfected Neuro2a cells with the targeting plasmid and observed high transfection efficiency based on mCherry signal (Fig. 2b). *Slc35a2* mRNA expression (Fig. 2c) and SLC35A2 protein expression (Fig. 2d) were significantly reduced in KO cells compared to wild-type and scramble controls.

**Fig. 2.**
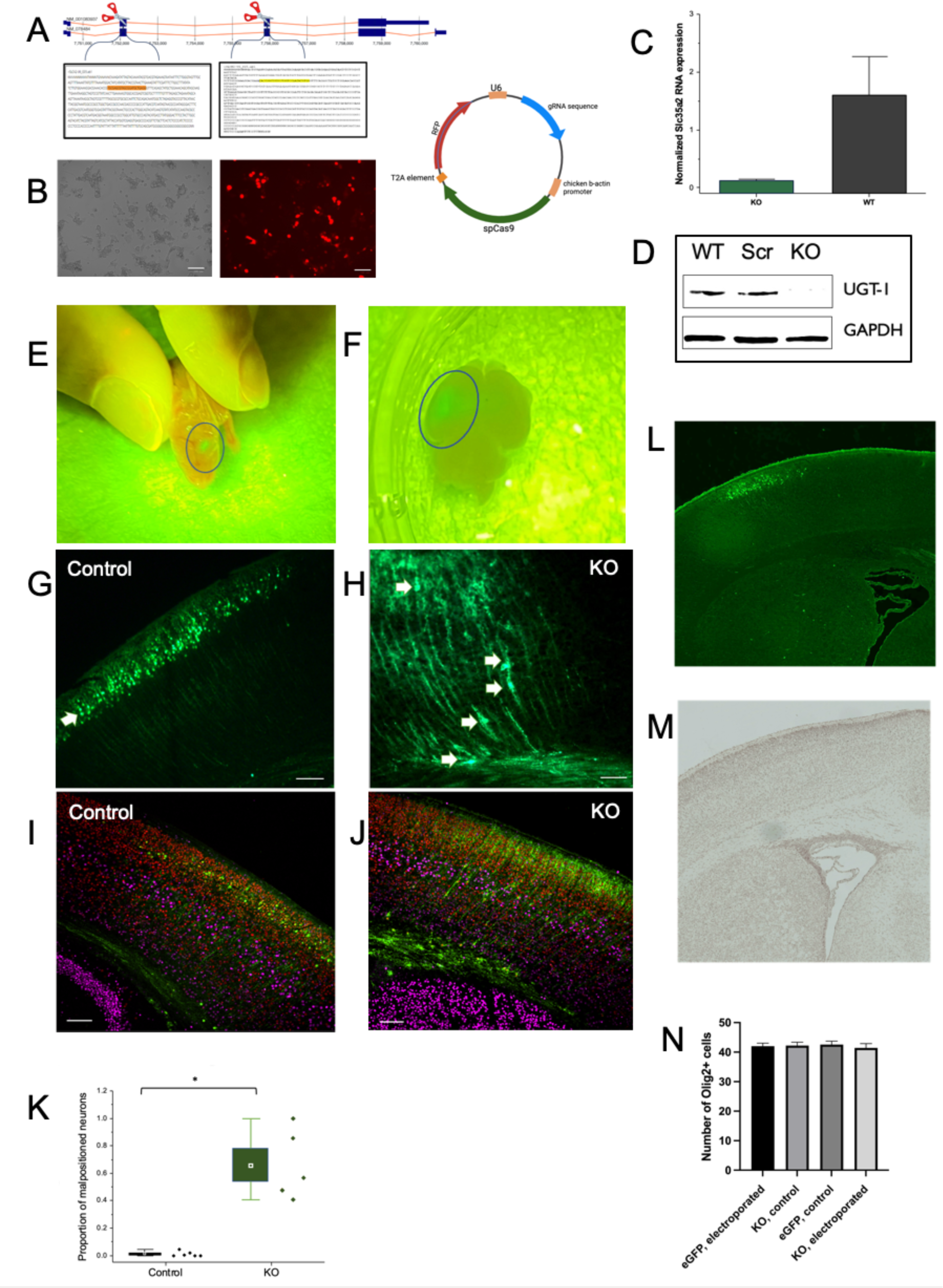
*Slc35a2* knockout results in altered cortical lamination. A) gRNAs targeting exons 2 & 3 spliced into pX458v2-R-spCas9 plasmid containing an mCherry fluorescent reporter. B) Neuro2A cells were transfected with double gRNA knockout (KO) plasmid and FACS sorted for mCherry fluorescence to create the Slc35a2 KO cell line. C) Slc35a2 KO cells have minimal Slc35a2 mRNA expression relative to GAPDH when compared to wild-type (WT) cells. D) KO cell lysates show no SLC35A2 expression compared to control lysates (scrambled gRNA sequence and WT). E) Slc35a2 KO mice at P0 show EGFP fluorescence indicating expression of KO plasmid in focal cortical area. F) Slc35a2 KO mice underwent transcardial perfusion and brain extraction at P4. G) EGFP+ control brain versus H) KO brain at P4 showing the distribution of positively transfected neurons (green). I) Control brain versus J) KO brain showing EGFP-positive cells in the context of their laminar destination with co-staining for cortical markers SATB2 (red, Layer II/III marker) and Ctip2 (magenta, Layer V marker). K) Proportion of malpositioned neurons (number of neurons not located in Layer II/III divided by total positively transfected neuron count) in Slc35a2 KO mice vs EGFP control transfected mice. L-M) GFP and Olig2 immunoreactivity in the cortex and white matter of a KO brain. (N) No significant difference in Olig2+ cell numbers were seen across groups. * P<0.05.

### *Slc35a2* knockout disrupts cortical lamination *in vivo*

To test the effect of *Slc35a2* KO *in vivo,* CRISPR gRNA-plasmids (or EGFP control plasmids) were delivered via *in utero* electroporation into the lateral ventricle of mice at E14 as a strategy to target cortical neurons destined for layer II/III. Pups were screened for EGFP-positivity (Fig. 2e) and brains were collected postnatally (Fig. 2f) to visualize EGFP-positive transfected neurons, as well as SATB2 and Ctip2 as laminar markers for cortical layers II/III and V, respectively. As expected, all EGFP-positive neurons transfected with the EGFP-only control plasmid were observed appropriately in layer II/III with none observed in deeper layers or sub-cortical white matter (Fig. 2g). In contrast, *Slc35a2* KO neurons were observed in cortical layers IV-VI, as well as within the subcortical white matter (n=6 animals per group, 3 sections per animal)(Fig. 2h). Few *Slc35a2* KO neurons were seen in layers II/III. EGFP-control neurons exhibited SATB2 expression (Fig. 2i), whereas only a subset of *Slc35a2* KO neurons expressed SATB2 (Fig. 2j). In both *Slc35a2* KO and EGFP control sections, Ctip2 expression identified neurons in an intact cortical layer V. None of the transfected neurons (*Slc35a2* KO or EGFP control) expressed Ctip2, which is consistent with a laminar birthdate destination of layer II/III following transfection on E14. Overall, significantly more (greater than 60%) transfected neurons in the KO condition were positioned outside of layer II/III compared to control (P<0.05; Fig. 2k).

### Oligodendrocyte density and cortical architecture

Given that MOGHE is characterized by oligodendroglial hyperplasia, we investigated whether focal *Slc35a2* KO would induce changes in oligodendrocyte cell density. We probed for Olig2 expression across the brain in *Slc35a2* KO, EGFP control, and WT mice such that 3 anatomical landmark sections were used per animal (n=3 animals per group, n=3 sections per animal): an anterior section displaying the lateral ventricle and fully formed corpus callosum, a section displaying the ventral hippocampal commissure and fimbria, and a section displaying the hippocampus. These sections allowed us to visualize within the electroporated hemisphere and unelectroporated hemisphere, with clear anatomical landmarks for congruency across samples. We observed no significant change in Olig2-positive cell count across groups (Fig. 2l-n). Given that IUE predominantly targets cortical neurons, we suggest that the oligodendroglial hyperplasia observed in MOGHE may result from a cell autonomous effect of *Slc35a2* loss in oligodendrocytes. This would be consistent with the observation that pathogenic *SLC35A2* variants are found in both neurons and oligodendrocytes in patient brain tissue^9^. Finally, to investigate gross cortical structure following IUE, we examined cresyl violet staining of coronal sections from *Slc35a2* KO, WT, and EGFP control mice. We observed no differences in overall brain structure, cortical lamination, or cortical thickness consistent with the sparse focal nature of the transfection (data not shown).

### *Slc35a2* knockdown is sufficient to disrupt cortical lamination

Some pathogenic *SLC35A2* variants linked to MOGHE may not completely abolish SLC35A2 protein function, though the effects have not yet been tested^10^. Some variant-containing cells from patients with *SLC35A2*-congenital disorder of glycosylation retain partial UDP-galactose transport activity^3^. Therefore, we asked whether *Slc35a2* KD and reduced Slc35a2 expression is sufficient to disrupt neuronal migration to an extent that is similar to *Slc35a2* KO.

Short-hairpin RNA (shRNA) sequences targeting *Slc35a2* were designed and cloned into plasmids co-expressing EGFP. Two independent shRNA sequences were used to verify the findings and results were compared to electroporations using a scrambled non-targeting shRNA sequence as a control. To test the efficiency of knockdown, we transfected cultured mouse NIH/3T3 cells with the constructs. Quantitative real-time PCR analysis of *Slc35a2* expression in transfected cells demonstrated knockdown efficiency of approximately 50% compared to a scrambled shRNA control in the context of ∼60% transfection efficiency (Fig. 3a).

**Fig. 3.**
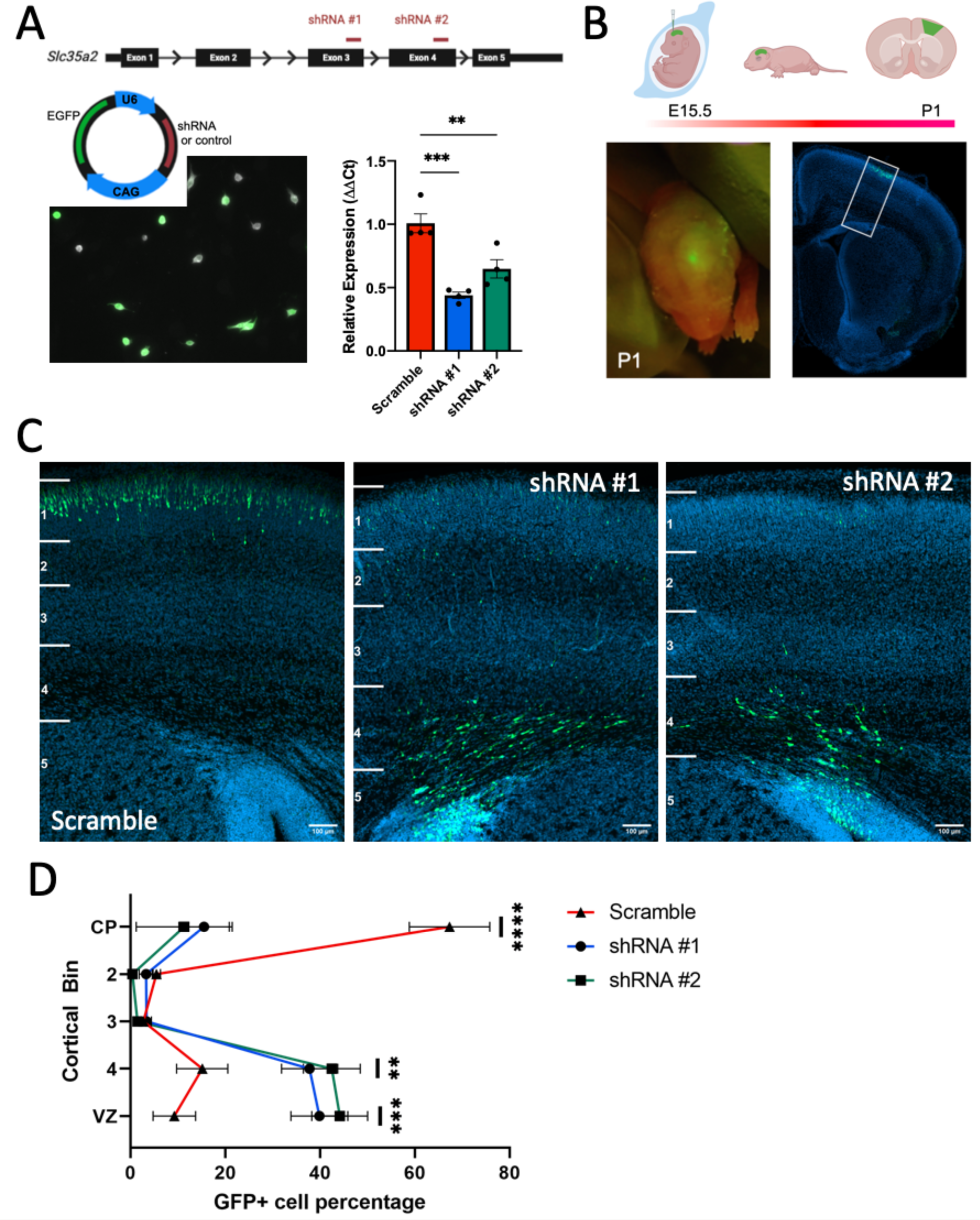
*Slc35a2* knockdown is sufficient to disrupt cortical lamination. A) shRNA sequences targeting *Slc35a2* or a scrambled control sequence were expressed in a U6-shRNA plasmid with EGFP as a reporter. Mouse NIH/3T3 cells were transfected with plasmid DNA to assess KD efficiency. *Slc35a2* mRNA expression was significantly reduced with shRNA treatment compared to scrambled control. B) Plasmid DNA was delivered to the lateral ventricle of E15.5 mouse embryos by IUE. C) Distribution of EGFP-positive cells in the mouse cortex at P1. D) KD resulted in significantly more neurons located in the deep cortical layers and subcortical white matter. Scale bars = 100 µm. Mean ± SEM. ** P<0.01, ** P<0.001.

We introduced the shRNA plasmids into the lateral ventricle of the embryonic mouse brain at E15.5 via *in utero* electroporation. At E15.5, neural progenitor cells lining the ventricle are destined to become layer II/III cortical neurons. Brains were collected and analyzed for EGFP expression at postnatal day 1 (Fig. 3b). We examined EGFP expression in coronal sections through the cortex and observed distinct profiles in knockdown versus control mice. EGFP-positive cells were mostly located in deep cortical layers and white matter in knockdown mice, whereas they were mainly restricted to outer cortical layers as expected in the control mice (Fig. 3c). We quantified the proportion of EGFP signal located in six bins spanning from the ventricular zone to the outer cortex, which revealed a significantly reduced percentage of cells properly located in the outer cortex of both groups of knockdown mice (Fig. 3d).

### *Slc35a2* KO lowers seizure threshold

To observe any behavioral or electrographic seizure activity following focal *Slc35a2* KO, mice were implanted with dural EEG electrodes and recorded for 72 hours post-recovery (Fig. 4a-c). No spontaneous seizures (behavioral or electrographic) were observed across any groups (WT, EGFP control or *Slc35a2* KO). We defined an electrographic seizure as 10 seconds or more of sharply contoured, rhythmic wave forms with a clear onset and clear offset associated with a Racine score of 3 or higher. Next, we assessed seizure threshold in mice by injecting mice with PTZ (55 mg/kg intraperitoneally) while under video and EEG monitoring^19^. Latency to first seizure as defined by electrographic activity was calculated from the exact time of injection. All electrographically defined seizures were also associated with behavioral seizures with a Racine score of 3-5. PTZ testing was conducted for EGFP control, WT, and *Slc35a2* KO mice at P60 and P120. At both timepoints, there was a significant difference in seizure latency between EGFP and *Slc35a2* KO groups as well as between WT and *Slc35a2* KO groups (p<0.05, unpaired t-test with Welch’s correction)(Fig. 4d-e). There was no difference between EGFP and WT groups at either age. These data demonstrate that focal *Slc35a2* KO increases sensitivity to PTZ, although it does not lead to spontaneous seizures in CD-1 mice.

**Fig. 4.**
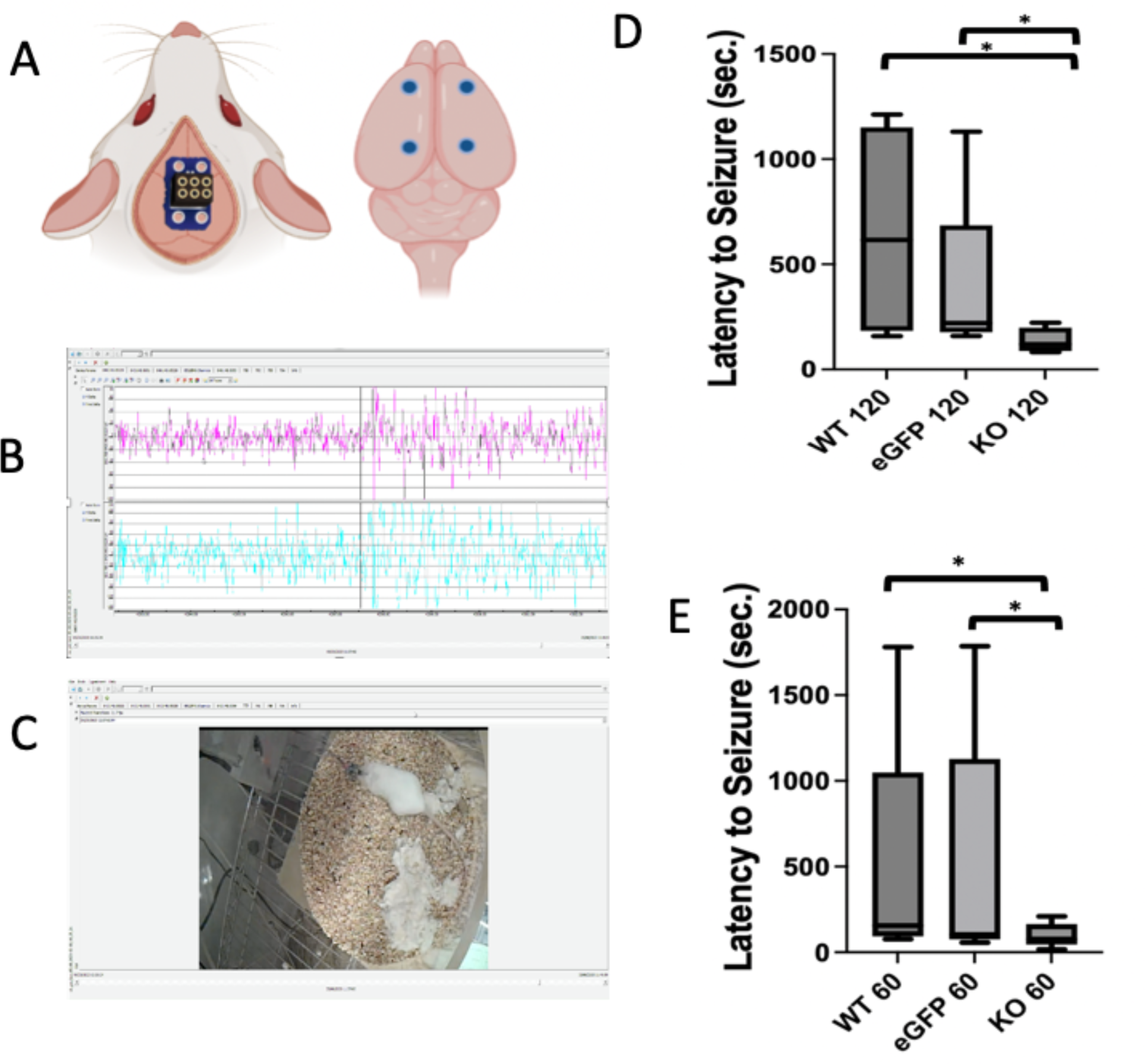
*Slc35a2* knockout results in a lower seizure threshold. A) Mice (*Slc35a2* KO, EGFP control, and WT littermates) were implanted with EEG electrodes. B,C) Video-EEG was recorded for 72 hours of baseline activity followed by PTZ threshold testing (55 mg/kg PTZ i.p). D) *Slc35a2* KO mice had a reduced seizure threshold when compared to both control groups at age P120 and at P60. *P<0.05.

## Discussion

Somatic *SLC35A2* variants represent an important cause of pediatric drug-resistant epilepsy and MCD. Here we demonstrate that either KO or KD of *Slc35a2* in mice leads to altered migration of transfected neurons during cortical development, heterotopic neurons identified in the subcortical white matter, and reduced seizure threshold in response to PTZ treatment. These data demonstrate that *Slc35a2* KO or KD leads to impaired laminar positioning of transfected neurons similar to histopathology observed in patient brain tissue. Heterotopic neurons are a consistent histopathological feature observed in both FCD I and MOGHE; our results support a direct role for SLC35A2 in neuronal migration and cortical lamination. Notably, *Slc35a2* KO in neurons via IUE did not induce changes in oligodendrocyte number, although oligodendroglial hyperplasia is a defining feature of MOGHE. This suggests that effects of *Slc35a2* loss in neurons and oligodendrocytes may result from distinct cell autonomous effects. Indeed, a recent study used a combined approach of laser capture microdissection and subsequent gene sequencing on human brain samples to identify *SLC35A2* variants in both heterotopic neurons and oligodendrocytes. ^9^

*SLC35A2* encodes the transmembrane protein SLC35A2, a transporter that shuttles UDP-galactose across the Golgi membrane. Deficient galactose transport impairs protein and lipid glycosylation, which has downstream effects on protein stability, localization, and transport. In addition to refractory epilepsy and intellectual disability, patients with brain somatic mosaic variants in this gene also face the added difficulty of a diagnosis. To date, patients have only been identified by analyzing surgically resected brain tissue samples, as the somatic variants in *SLC35A2* are only expressed in focal regions of the brain. The pathogenic variants in *SLC35A2* are varied in type (i.e., missense, frameshift, nonsense), frequency and distribution throughout the affected tissues, as well as localization within the protein itself (any of the 10 predicted transmembrane domains or cytosolic loops)^7^. Higher variant allele fraction is linked to more extensive histopathological disorganization and epileptic activity^11^. Recent studies provide insights into genotype-phenotype correlations based on the location of variants in the protein domains as well as the variant allele fraction per patient.^10^ Our experiments add to this literature by showing sparse focal KO or KD of *Slc35a2* in mice is sufficient to alter cortical development.

While *Slc35a2* KO was not sufficient to cause spontaneous seizures, it did result in a reduced seizure threshold in response to PTZ. These findings demonstrate a mechanistic link between focal loss of *Slc35a2* and enhanced seizure susceptibility. Our data show that reduced SLC35A2 function has profound effects on assembly of the neocortex which may result in altered brain circuitry leading to seizure susceptibility. Recent studies have shown that patients with SLC35A2-CDG who receive oral D-galactose supplementation show improvements in cognition and reduced seizure frequency^20^. These trials are straightforward in patients with germline *SLC35A2* variants that can be identified easily versus somatic conditions like FCD I or MOGHE where brain tissue is required to detect the *SLC35A2* variant. Future studies to assess the role of D-galactose supplementation will be important in patients harboring somatic variants in *SLC35A2*.

In summary, these experiments are the first to provide a direct experimental link between *Slc35a2* and cortical development in mice. Our results demonstrate that neuronal migration deficits are likely a key contributor to the MCD observed in patients with somatic *SLC35A2* variants. Future studies will further investigate the mechanisms and potential treatment opportunities associated with *SLC35A2*-related epilepsies.

## Acknowledgements

The work was funded by NIH-NINDS (R01NS115017) to ELH/PC and NIH-NINDS (R01NS129784) to TAB. The content is solely the responsibility of the authors and does not necessarily represent the official views of the NIH. Some figures were created in BioRender.

## Author Contributions

SE, SS, HY, RC, JP, AR, SL, JF, AL and AR performed the experiments. SE and SS analyzed the results. ELH and PI provided resources and input. PC and TAB supervised the research. SE, SS, PC and TAB wrote the original draft. All authors participated in review and editing of the manuscript.

